# Biofuel-driven Adaptable n-type Supramolecular Wires: Mimicking Conducting Microbial Nano Filaments

**DOI:** 10.1101/2025.11.12.687965

**Authors:** Sunil Kumar, Rohit Kapila, Arpita Panda, Pooja Singh, Veerbhadrarao Kaliginedi, Subinoy Rana

## Abstract

We report the biofuel-driven self-assembly of a benzimidazole–pyridine platinum (II) complex into supramolecular nanowires. Phosphate promotes Pt···Pt metallophilic stacking, whereas the pristine complex forms nanosheets dominated by π–π interactions. By varying phosphate denticity (adenosine/guanosine mono-, di-, and triphosphates), we tune the extent of Pt···Pt interactions and the resulting electrical conductance of the nanowires, measured on graphene using a custom EGaIn setup. Triphosphate-templated nanowires exhibit nearly four orders of magnitude higher conductance than non-templated nanosheets. Thermopower measurements reveal a negative Seebeck coefficient, indicating LUMO-mediated electron transport and rare n-type behavior in these supramolecular assemblies. Temperature-dependent measurements show a transition from tunneling (non-templated nanosheets) to thermally activated hopping (triphosphate templated nanowires). The system is recyclable, apyrase hydrolysis disrupts the nanowires, and re-addition of phosphate restores assembly, enabling reversible conductance switching over multiple cycles. This study demonstrates a fuel-responsive, adaptive supramolecular electronic material, highlighting how controlled Pt···Pt interactions can be exploited to program charge transport in soft, bioinspired electronic systems.

## INTRODUCTION

Biological systems have long inspired the design of synthetic materials with advanced functionalities, particularly in the realm of long-range electron transport.^1–6^ Microbial nanowires, particularly those composed of multi-heme cytochromes like OmcS and OmcZ, have emerged as promising bioelectronic materials due to their unique electron transport properties.^7–9^ Studies have shown that temperature-induced restructuring of hydrogen bonds can enhance their conductivity by several hundred-fold.^10^ Structural insights into OmcZ nanowires revealed densely packed hemes that facilitate efficient electron hopping,^8^ while synthetic protein nanowires now offer programmable assembly and tunable functionality. Despite the central role of OmcS in some strains, alternative extracellular electron transfer (EET) routes, including conductive pilli, have also been reported.^11^ These discoveries highlight the dynamic, diverse mechanisms behind long-range electron transport in microbial systems and their potential in bioelectronic applications.^12–15^ The exceptional conductivity of microbial nanowires arises from precise molecular templating and non-covalent interactions, enabling organization that significantly outperforms non-heme filaments.^16^ Mimicking these natural conductors is highly attractive, as they offer a rare combination of efficiency, adaptability, and environmental sustainability—traits often lacking in conventional electronic materials. Synthetic supramolecular electronics seeks to emulate such efficiency by harnessing non-covalent forces such as π-π stacking, metal-metal interactions, and hydrogen bonding to assemble molecular units into functional nanoscale systems.^17^ However, replicating the structural complexity and functional versatility of proteins like OmcS poses significant challenges, necessitating innovative approaches in synthetic systems.

Recent breakthroughs in supramolecular science have enabled the creation of adaptable materials emulating different biological systems including proteins, enzymes, cytoskeleton, cellular organelles, and minimal cells.^18–22^ Adaptability is a hallmark of living systems that incorporates energy consuming processes to shuttle between assembly states, leading to varied stimuli responsiveness and functions. Artificial molecular machines that mimic biological motors by converting chemical energy into directional nanoscale motion, mark a major advance in synthetic molecular engineering^23–25^. Likewise, energy dissipating non-equilibrium assemblies have been introduced to fabricate supramolecular polymers, nanoreactors, coacervates, and membranes with spatiotemporal control of structure and function. Recent advances have demonstrated that synthetic assemblies, particularly those involving transition metal complexes, can emulate the charge transport capabilities of biological systems like proteins, while offering tunable properties for applications in sensors, diagnostics, and energy devices (Li et al., 2023 and others).^26,27^ Despite of availing these applications, n-type supramolecular assemblies—a rare case due to the easy trapping of electrons by O_2_/H_2_O under ambient conditions—add a tremendous importance for allowing these device fabrications based on thermoelectric applications where these assemblies reduce the material imbalance.^28–31^ These assemblies also provide balanced n/p thermoelectric legs for high thermoelectric performance, new pathways to electron-transport based optoelectronic biomedical devices.

Our prior work introduced self-assembly of a platinum(II) complex (**Pt-1**, Figure 1a) into nanosheet-like structures with aggregation-induced emissive and oxidase enzyme-like functions under light activation.^32^ We transformed these thermodynamically driven structures into non-equilibrium supramolecular assemblies using a multivalent fuel, adenosine triphosphate (ATP). The nature and extent of the supramolecular structure is governed by the biological fuel, which templates **Pt-1** along the backbone and modulates the extent of Pt···Pt and π-π interactions. This approach draws parallels to the heme-stacking mechanism in microbial nanowires, where precise molecular organization amplifies electron transport, and extends it into the realm of out-of-equilibrium synthetic systems.^33^ Our experimental findings using widely adopted EGaIn junction technique^34^ (figure 1b and 1c) demonstrate that ATP-templated **Pt-1** nanowire structures exhibit conductance enhancement exceeding three orders of magnitude compared to the non-templated complex **Pt-1**, suggesting a critical role of Pt···Pt interactions in governing the charge transport through these supramolecular nanowires. This study investigates a critical question on how fuel-driven metal-metal interactions modulate the structural and electronic properties of supramolecular assemblies. Further, the transient assembly state of **Pt-1** achieved by an enzyme-controlled negative feedback loop shows promise in harnessing these nanowires for recyclable, bioinspired electronic materials.

**Figure 1.**
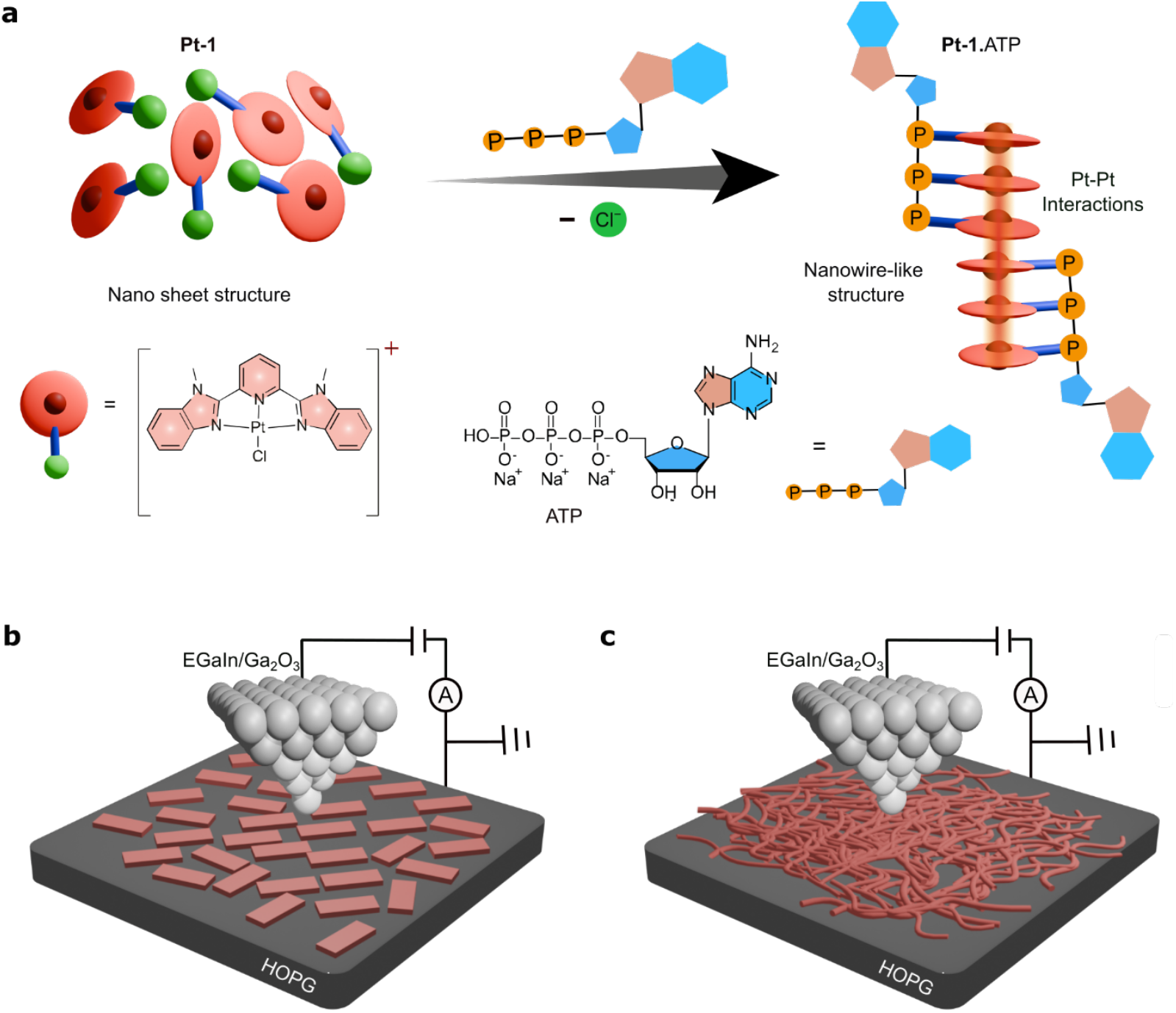
Strategy for Assembly Formation and Electrical Probing. **a**, Schematic illustration of the structural transformation from nanosheet-like Pt-1 complex to nanowire-like **Pt-1**.ATP assembly induced by the biofuel ATP. In the **Pt-1** complex, a platinum centre (red sphere) forms a coordination bond with a conjugated ligand environment (light red disc) and a covalent bond with a chloride ion (green sphere attached via a blue line). Upon ATP addition, the chloride is replaced by a phosphate group (yellow sphere), promoting π-stacking and enhanced Pt–Pt interactions (indicated by overlapping yellow and red shades), which drives the reorganization into nanowire morphology. **b, c**, Schematic diagrams of the electrical measurement setup. A eutectic gallium–indium (EGaIn) alloy tip (74.6% Ga, 24.6% In) is used as the top electrode to probe the assemblies of **Pt-1 (b)** and **Pt-1**.ATP **(c)** spin coated on a grounded HOPG (highly ordered pyrolytic graphite) substrate.

## RESULTS AND DISCUSSION

### Fabrication of self-assembled metallo-supramolecular wires: Fabrication of self-assembled metallo-supramolecular wires

To construct supramolecular nanowires with tunable coordination dynamics, we designed and synthesised a methyl-substituted benzimidazole–pyridine pincer ligand (mBzPy) that stabilises a platinum(II) centre, affording the complex **Pt–1** (Figure 1a). The complex was obtained via direct reaction between K_2_PtCl_4_ and the mBzPy ligand, yielding a mononuclear Pt(II) species bearing a labile chloro ligand. We anticipated that ionizable chloride facilitates displacement by biologically relevant triphosphate anions such as ATP, forming a **Pt-1**.ATP conjugate through coordination between the Pt(II)centre and the phosphate oxyanions. This conjugate serves as a molecular scaffold that directs the hierarchical self-assembly of supramolecular nanowires via synergistic π-π stacking and Pt···Pt metallophilic interactions. For comparison, a *σ*-alkynyl platinum(II) complex (**Pt–2**) lacking a replaceable chloride ligand was also synthesized, which exhibited only electrostatic association with ATP. Both complexes were isolated in moderate to good yields (Supplementary Scheme S1– S4) and were fully characterized by ^1^H and ^13^C NMR spectroscopy and mass spectrometry (Supplementary Figures S1–S12).

To probe the coordination-driven assembly, we performed ^1^H NMR titration experiments of **Pt-1** with ATP, which revealed a progressive downfield shift of proton resonance, indicative of strong interactions between the electronegative phosphate oxyanions of ATP and the Pt(II) centre (Figure 2a). For additional solution characterizations, we monitored the UV-vis spectra of **Pt-1** upon the addition of ATP. The **Pt-1**.ATP complex displayed an enhanced band at 550 nm, which is attributed to metal-to-metal-ligand charge transfer (MMLCT) (Figure 2b). This ^1^MMLCT band is attributed to enhanced noncovalent interactions such as π-π and Pt···Pt interaction upon assembly formation.^35^ Similarly, the photoluminescence (PL) corresponding to the MMLCT band of **Pt-1** amplified with increasing ATP concentration upon excitation at 390 nm (Figure 2c). Further, the binding mode of templated self-assembly of **Pt-1** with ATP by replacing the chloro group was established through X-ray photoelectron spectroscopy (XPS) studies. XPS analysis revealed a difference in the electronic properties of platinum centre in the assemblies on a highly oriented pyrolytic graphene (HOPG) substrate. The Pt 4f_7/2_ spectrum of **Pt-1** showed a binding energy of 72.3 eV that shifted to a higher binding energy of 72.9 eV upon **Pt-1**.ATP adduct formation (Figure 2d), suggesting more interaction between platinum and the oxyanions of ATP in the assembly. The XPS analyses also provided insights into the Pt–Cl bond, which shows binding energies of 198.4 eV for Cl 2p_3/2_ and 200.1 eV for Cl 2p_1/2_ (Figure S13-S14). These values decrease to 197.6 eV and 199.3 eV, respectively, upon formation of **Pt-1**.ATP adducts, indicating a weakening of the Pt–Cl bond due to the ATP binding. In contrast, the XPS peaks of carbon and nitrogen atoms remain unchanged (Figure S13-14). Raman spectroscopy further supports the presence of Pt···Pt interactions as indicated by the increase in the intensity of the vibrational band in the range of 100 to 220 cm^-1^ (Figure 2e). Altogether, these studies confirm ATP-induced displacement of the chloro group of **Pt-1**. To gain further insight into morphological variations, the formed assemblies were examined using tapping-mode atomic force microscopy (AFM) on thin films. The **Pt-1** complex alone displayed short-range self-assemblies in a DMSO:water (1:1 v/v) solvent mixture, exhibiting a discrete nanosheet morphology with a length of 200-300 nm and a thickness of ∼1-2 nm on a HOPG substrate (Figure 2f). Notably, under similar conditions, the interaction between the **Pt-1** and ATP resulted in the formation of interconnected nanowires of >10 μm length with a similar height profile, which were homogeneously distributed across the HOPG surface (Figure 2g). These morphological changes further confirm the templated hierarchical assembly of **Pt-1**, resulting from π-π and Pt···Pt interactions.

**Figure 2.**
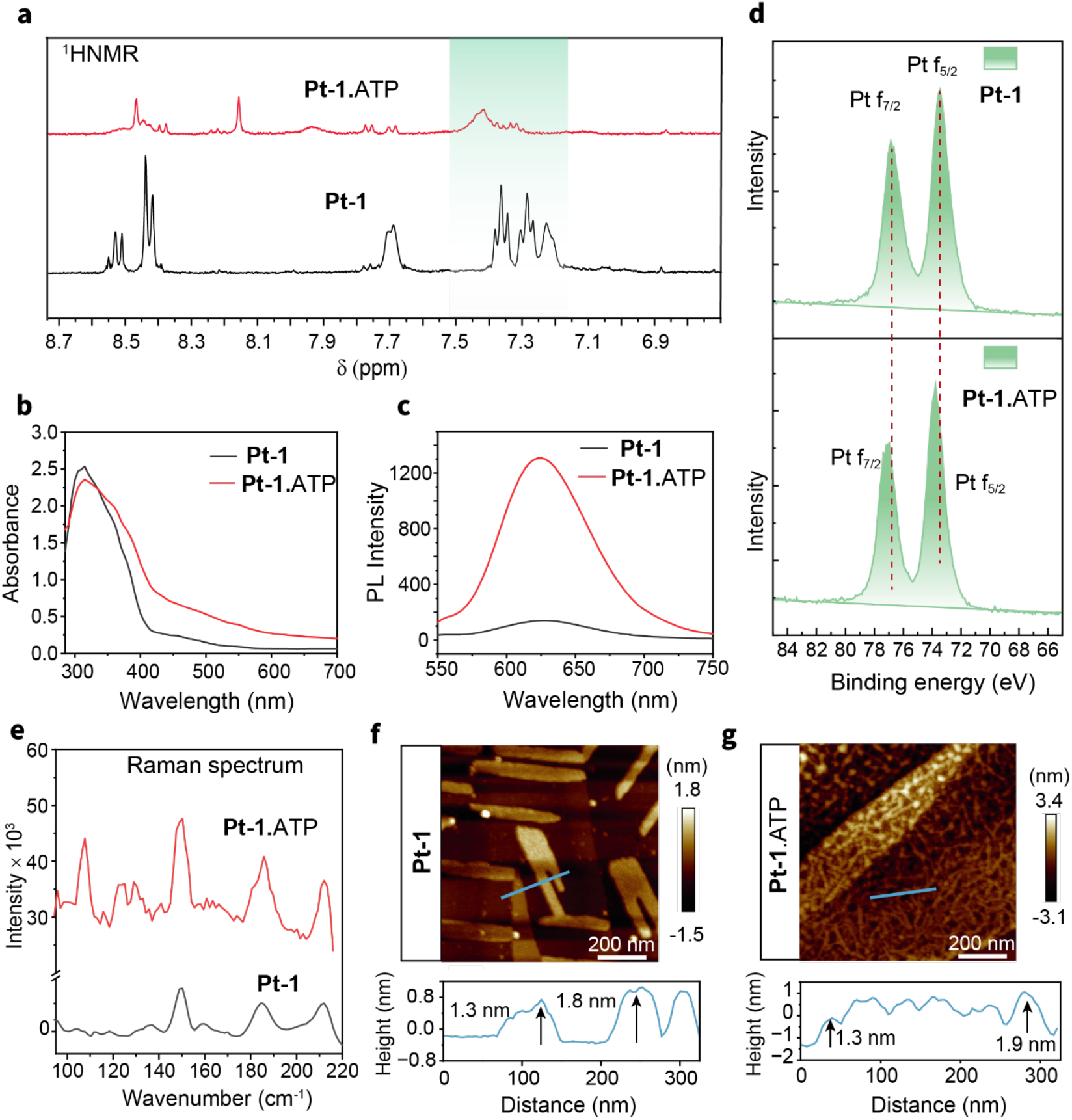
Structural and spectroscopic characterization of Pt-1 and Pt-1.ATP assemblies. **a**,^1^H NMR spectra of **Pt-1** (black) and **Pt-1**.ATP (red), showing the downfield shift of peaks upon addition of ATP, indicative of strong interactions of phosphates. **b**, UV–vis absorption spectra of **Pt-1** and **Pt-1**.ATP, showing enhanced absorption near 550 nm in the ATP-bound state. **c**, Photoluminescence (PL) spectra of **Pt-1** and **Pt-1**.ATP, exhibiting increased emission intensity upon ATP addition. **d**, X-ray photoelectron spectra (XPS) of **Pt-1** and **Pt-1**.ATP, showing shifts in Pt 4f_7/2_ and 4f_5/2_ binding energies (red dotted lines), consistent with increased electronic interaction between platinum centres and the phosphate groups of ATP. **e**, Raman spectra of **Pt-1** (black) and **Pt-1**.ATP (red), displaying the emergence of a peak at 110 cm^-1,^ corresponds to strong Pt···Pt interactions upon ATP binding. **f**, Atomic force microscopy (AFM) image of **Pt-1** over a 1 µm × 1 µm area, showing nanosheet morphology. **g**, AFM image of **Pt-1**.ATP over the same area, revealing long nanowire structures. Corresponding height profiles (below) confirm the presence of consistent nanoscale features.

### Electrical characteristics of the supramolecular assemblies

Given the long-range ordering of the Pt(II) centres induced by templation, we examined the electrical conductance of the templated assemblies on HOPG substrate. The electrical characterization revealed a striking difference in current density between the **Pt-1**.ATP adduct and pristine **Pt-1** complex. We observed a nearly four orders of magnitude enhancement in ATP-templated assembly compared to the **Pt-1** (Figure 3a). This substantial enhancement underscores the profound effect of Pt···Pt metallophilic interactions on the electronic properties of the assembly upon ATP templation. In contrast, the short assemblies of **Pt-1** complex do not provide significant Pt···Pt interactions, resulting in much lower conductivity. The difference in electrical behavior can be attributed to the distinct morphological and structural features of the assemblies—the **Pt-1**.ATP adduct forms an extended nanowire network on the substrate, as observed in the morphological analysis (Figure 2g). These findings highlight the critical role of ATP in modulating the structure and electronic properties of supramolecular assemblies, showcasing the potential of such systems for tunable and high-performance electronic applications.^26,36^

**Figure 3.**
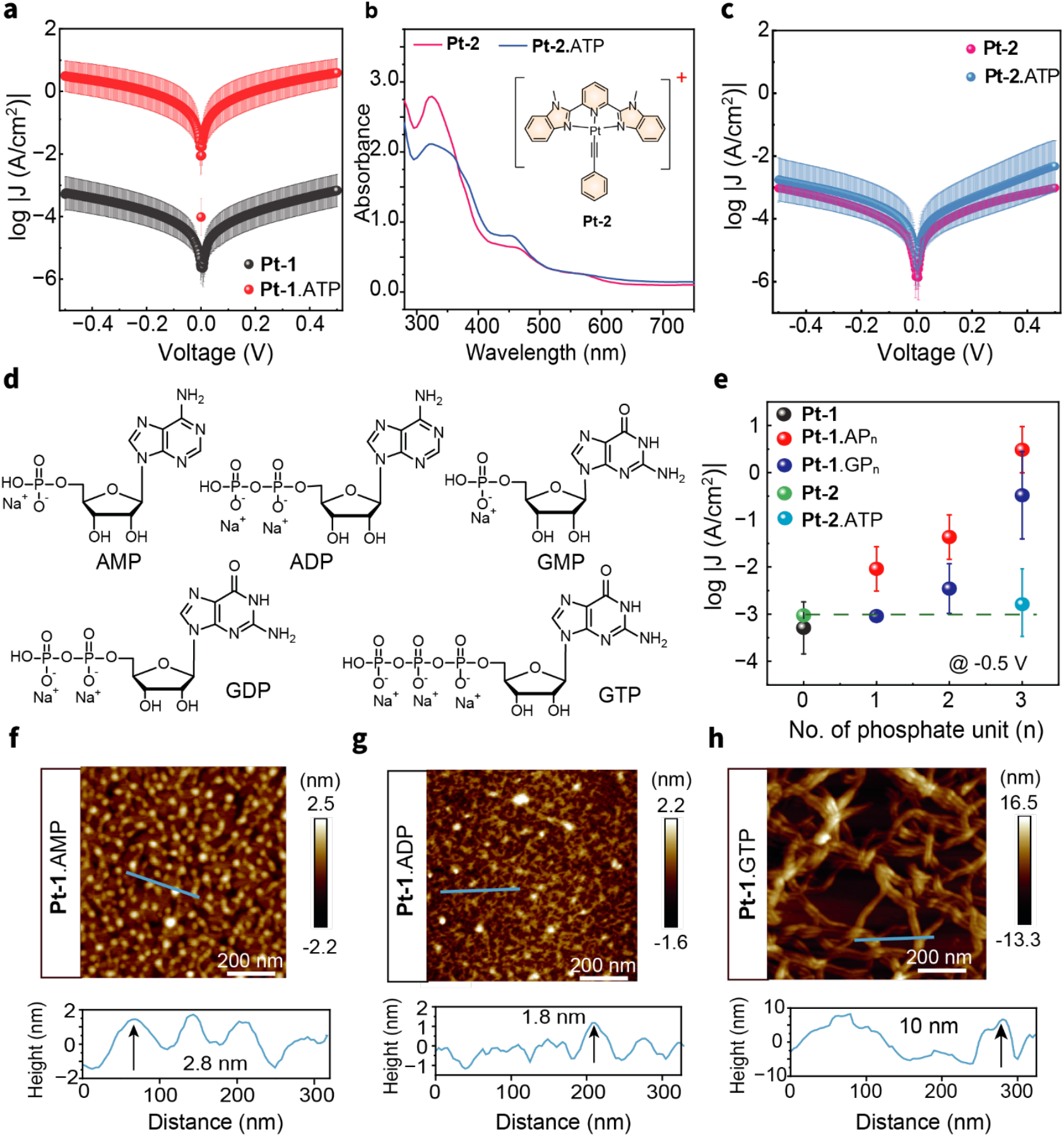
Electrical and morphological responses of Pt complexes upon interaction with phosphate-containing molecules. **a**, Plot of log|J| versus voltage (V) for **Pt-1** (black) and **Pt-1**.ATP (red) in the range of – 0.5 V to +0.5 V, showing a significant enhancement of current density, approximately four orders of magnitude higher for the **Pt-1**.ATP adduct compared to the pristine **Pt-1** complex, highlighting ATP-induced electronic modulation. **b**, UV-visible spectra of **Pt-2** (pink) and **Pt-2**.ATP (blue) along with the molecular structure of **Pt-2**. Minimal change in ^1^MMLCT at 550 nm upon ATP addition, attributed to the absence of Pt···Pt interactions, which are suppressed by introducing a sterically rigid and non-replacable unit in place of the labile chloride (Cl^−^) ligand. **c**, Plot of log|J| versus voltage for **Pt-2** (pink) and **Pt-2**.ATP (blue), recorded from –0.5 V to +0.5 V, revealing negligible change in current density, further supporting the suppression of ATP-mediated interaction in this system. **d**, Molecular structures of selected phosphate-containing biomolecules—guanosine monophosphate (GMP), adenosine monophosphate (AMP), adenosine diphosphate (ADP), guanosine diphosphate (GDP), and guanosine triphosphate (GTP)used to study phosphate unit dependence in electrical response. **e**, Plot of log|J| as a function of the number of phosphate units [n =1 (monophosphate), 2 (diphosphate) and 3 (triphosphate)] showing a gradual increase in current density with increasing phosphate content, including comparative data for **Pt-2** and **Pt-2**.ATP assemblies. **f–h**, atomic force microscopy (AFM) images (1 µm × 1 µm) of **Pt-1**.AMP **(f), Pt-1**.ADP **(g)**, and **Pt-1**.GTP **(h)**, illustrating the evolution of nanostructures upon interaction with different phosphate-containing molecules. The height profiles below each image indicate similar feature dimensions for **Pt-1**.AMP and **Pt-1**.ADP, whereas **Pt-1**.GTP exhibits broader structures, with vertical features ranging from ∼8 to 10 nm. AFM data for **Pt-1**.GMP and **Pt-1**.

To validate the role of Pt-phosphate coordination in directing the supramolecular nanowires, we studied a control complex (**Pt-2)**, which lacks the labile chloro group. Unlike **Pt-1**, the electrostatic interaction between **Pt-2** and ATP under similar conditions showed a negligible change in the ^1^MMLCT band at 550 nm in the UV-Vis spectrum, indicating the absence of Pt···Pt interactions (Figure 3b). Furthermore, the PL spectra corresponding to the electrostatic assembly showed a negligible change relative to the pristine **Pt-2** complex (Figure S15). Electrical measurements of the **Pt-2** complex and its ATP adduct revealed comparable current densities within the same order of magnitude (Figure 3c). This finding contrasts sharply with the pronounced conductance enhancement observed for the **Pt-1**.ATP assembly, underscoring the importance of Pt-phosphate coordination in promoting long-range electronic coupling. The rigid Pt–C≡C–Ph bond in **Pt-2** prevents the formation of ordered arrangements upon ATP addition, restricting interactions to electrostatic associations rather than the coordination-driven templation seen in **Pt-1**.ATP. As a result, the **Pt-2** assemblies remain structurally disordered, yielding smaller aggregates with minimal Pt···Pt interactions, even in the presence of ATP. Morphological analyses further corroborate these observations, revealing the absence of extended nanowire formation like **Pt-1**.ATP system. The **Pt-2** complex exhibited a dotted cluster-like morphology (Figure S16), distinct from the long nanowire structures observed in the **Pt-1**.ATP assembly. When ATP was introduced, the **Pt-2**.ATP adduct formed flakes and hollow, faceted, rod-like structures (Figure S17). Collectively, these results emphasise the role of the labile chloro ligand in **Pt-1**, which enables coordinative templation with ATP to drive long-range Pt···Pt coupling and the formation of highly ordered and conductive supramolecular nanowires.

Building upon the pivotal role of the labile chloro ligand in facilitating ATP coordination, we next explored how the denticity of phosphate templates influences the assembly and electronic behaviour of **Pt-1** complex. To elucidate this effect, we extended our electrical measurements to a series of phosphate-containing nucleotides, namely adenosine monophosphate (AMP), adenosine diphosphate (ADP), guanosine monophosphate (GMP), guanosine diphosphate (GDP), and guanosine triphosphate (GTP) (Figure 3d). This comparative analysis aimed to evaluate the selectivity and generalizability of our system toward other nucleotides with varying numbers of phosphate units. Upon morphological analysis, we observed the variation in the interconnectivity of the nanowire with an increase in the number of phosphate units evident across **Pt-1**.AMP, **Pt-1**.ADP (Figure 3f, 3g and S18-19), **Pt-1**.GMP, **Pt-1**.GDP and **Pt-1**.GTP (Figure S20-S22, and 3h, respectively). The electrical measurements followed a consistent trend, with current density increasing proportionally to the phosphate denticity in both adenosine and guanosine series (Figure 3e). Notably, the adenosine-derived assemblies exhibited approximately one order of magnitude higher conductance than their guanosine counterparts, likely due to stronger secondary interactions, including intermolecular hydrogen bonding and enhanced π–π stacking..^37,38^

In contrast, AMP and GMP, which contain only a single phosphate group, failed to produce similar structural or electrical enhancements (Figure S23a and S31a). These nucleotides exhibited weaker binding to the **Pt-1** complex compared to ATP and GTP. The **Pt-1**.AMP adduct displayed a modest, one order increase in current density over pristine **Pt-1** complex, whereas the **Pt-1**.GMP adduct showed a slight increase in current density compared to the **Pt-1** complex, although it was still lower than that of **Pt-1**.AMP (Figure 3e). These findings suggest that the assembly-enhancing effect observed with ATP is not specific to adenosine base pairs but rather a general characteristic of trivalent phosphate-driven assembly. The ability of GTP to induce similar structural and functional properties confirms that the triphosphate unit plays a central role in organizing the **Pt-1** complex into extended conductive nanowire structures. This insight broadens the scope of potential nucleotides or analogues that could be employed to modulate the properties of such supramolecular systems, paving the way for diverse applications in bioelectronics and nanotechnology.

### Determining the nature of charge carriers and mechanism of charge transport

The introduction of triphosphate fuel into the **Pt-1** complex assemblies, enhancing Pt-Pt interactions, raised intriguing questions about the nature of charge carriers and transport mechanism within these systems. To understand more about the charge carriers and transport process, we conducted thermopower measurements. We modified our measurement setup by employing a temperature sensor onto the substrate and a ceramic heater below the substrate controlled by temperature controller. The substrate was heated from the bottom side of the substrate, and the temperature was monitored by temperature sensor placed on top (Figure 4a). Thermopower is a measure of thermoelectric voltage of a material induced by temperature gradient.^39^

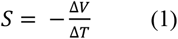

**Figure 4.**
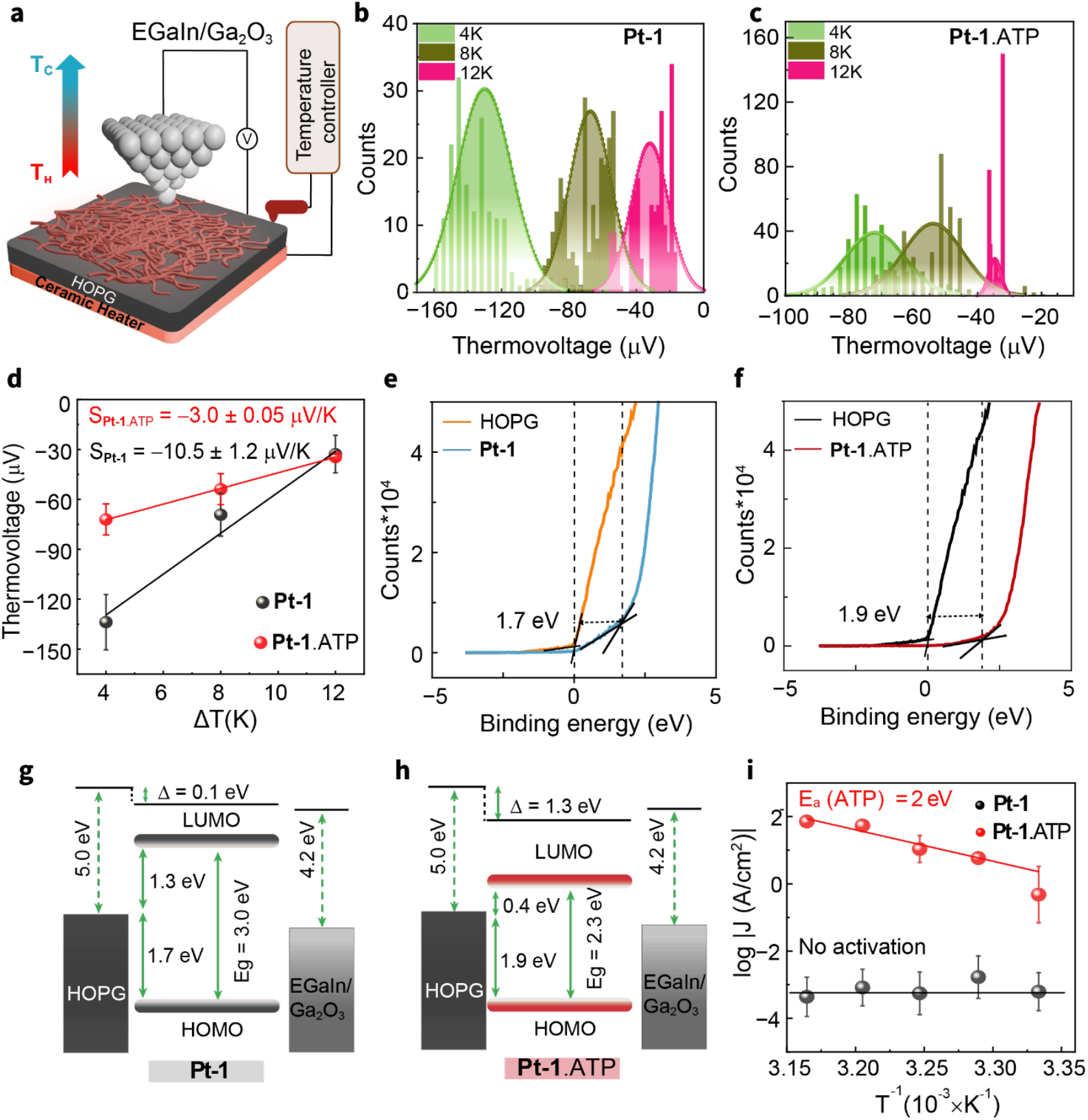
Thermoelectric and electronic characterization of Pt-1 and Pt-1.ATP assemblies. **a**, Schematic representation of the thermal measurement setup used for thermopower and temperature-dependent current– voltage (I–V) characterization. A ceramic heater is integrated beneath the substrate and controlled in a feedback loop with an external temperature sensor placed above the device, enabling the generation of a vertical temperature gradient across the substrate for Seebeck coefficient measurements. **b, c**, Histograms of measured thermovoltage (ΔV) at temperature differences (ΔT) of 4 K, 8 K, and 12 K for **Pt-1** (**b**) and **Pt-1**.ATP (**c**), each fitted with Gaussian distributions. **d**, Plot of thermovoltage (ΔV) as a function of temperature difference (ΔT) for **Pt-1** and **Pt-1**.ATP, showing a linear increase with positive slope, indicating negative Seebeck coefficients in both systems. This confirms electrons as the dominant charge carriers in the transport process. **e**, Ultraviolet photoelectron spectroscopy (UPS) spectra of **Pt-1** and **f**, Corresponding energy level diagram of EGaIn/Ga_2_O_3_//**Pt-1**/HOPG junction. **g**, Ultraviolet photoelectron spectroscopy (UPS) spectra of **Pt-1**.ATP and **h**, Corresponding energy level diagram of EGaIn/Ga_2_O_3_//**Pt-1**.ATP/HOPG junction, providing information on the HOMO energy level offset (E_F_ − E_HOMO_). Energy-level diagrams of **Pt-1** and **Pt-1**.ATP are constructed using the HOMO energy level offset values from UPS and the optical bandgap estimated from UV–vis absorption spectroscopy. A shift in energy alignment upon ATP binding is observed, indicating modified interfacial energetics. **i**, Arrhenius plots of log|J| versus inverse temperature (1/T, in 10^-3^ K^−1^) for **Pt-1** and **Pt-1**.ATP. The **Pt-1**.ATP assembly exhibits thermally activated transport behavior, whereas **Pt-1** shows negligible temperature dependence, consistent with a coherent tunneling transport mechanism.

Here, S represents the thermopower, also referred to as the Seebeck coefficient, measured in µV/K, while ΔT denotes the temperature gradient, and ΔV is the induced thermovoltage.

The Seebeck coefficient offers insights into the molecular orbitals involved in charge transport. A positive Seebeck coefficient indicates that charge transport is dominated by the HOMO (highest occupied molecular orbital), with holes serving as the primary charge carriers, whereas a negative Seebeck coefficient suggests transport dominated by the LUMO (lowest unoccupied molecular orbital), with electrons as the charge carriers. In our experiments, we measured temperature induced thermovoltage for three temperature differences of 4 K, 8 K, and 12 K under low humidity (Figure S32). We built gaussian fitted histograms for thermovoltage at each temperature difference for **Pt-1** and **Pt-1**.ATP (Figure 4b, 4c).The linear fit of these thermovoltage to the temperature difference ΔT results in a positive slope which indicates a negative Seebeck coefficient for both the **Pt-1** complex and **Pt-1**.ATP assemblies –10.5 ±1.2 µV/K and –3.2 ± 0.1 µV/K respectively (Figure 4d). This observation indicates that charge transport in these assemblies is LUMO-dominated, with electrons serving as the primary charge carriers.

To validate our findings from thermopower measurements, we performed Ultraviolet Photoelectron Spectroscopy (UPS) to find position of the HOMO in the **Pt-1** complex and **Pt-1**.ATP. The HOMO energy offset (E_F,HOMO_) for the **Pt-1** complex was determined to be at 1.7 eV, while for **Pt-1**.ATP, the HOMO onset was farther at approximately 1.9 eV (Figure 4e, 4f). The LUMO position was estimated by subtracting the (E_F,HOMO_) from the optical bandgap, calculated as 3.01eV for **Pt-1** and 2.32 eV for **Pt-1**.ATP using Tauc plots (Figure S33). The (E_F,LUMO_) for the **Pt-1** complex was found to be around 1.3 eV (Figure 4g), whereas for **Pt-1**.ATP, the LUMO lies much closer to the Fermi energy, at just 0.4 eV (Figure 4h). These observations solidify that electrons are the primary charge carriers in both assemblies. The significantly lower LUMO position in **Pt-1**.ATP explains its three-fold higher conductance compared to the bare **Pt-1** complex. UPS measurements also revealed that the work function of the **Pt-1** complex is almost identical to the HOPG substrate, with a vacuum shift of Δ=0.1eV. However, for **Pt-1**.ATP, a substantial vacuum shift of Δ = 1.3 eV was observed, indicating a significant decrease in work function, further supporting the enhanced conductance in this system.

To understand the charge transport mechanism in these assemblies, we conducted temperature-dependent I-V measurements^40^. A total of 220–250 I-V curves (forward and backward) were recorded across five temperatures: 27°C, 31°C, 35°C, 39°C, and 43°C (substrate temperature) (Figure S34-S35). To discriminate the temperature dependence of current density, we fitted our data with Arrhenius equation:

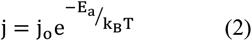

Where k_B_ is Boltzmann constant (*k*_*B*_ = 8.62 × 10-5 eV/K), j_o_ is pre-exponential factor and E_a_ is the activation energy. The results shown in (Figure 4i), reveal that the current density for the **Pt-1** complex remains unaffected by temperature, indicating temperature-independent coherent tunneling. In contrast, **Pt-1**.ATP assembly exhibits thermally activated current density changes, following the Arrhenius model, with activation energy close to ∼2 eV. A similar magnitude of activation energy was observed with **Pt-1**.GTP assembly (Figure S35,f). This high activation energy, compared to similar molecular systems of the same thickness, underscores the significant temperature sensitivity of these fiber structures.^41–43^ These findings confirm that while charge transport in the **Pt-1** complex is dominated by coherent tunneling, the introduction of triphosphate fuel enhances both the current density and temperature dependence of the assemblies.

### Transient nature of supramolecular assembly and Conductivity

Adaptable structure regulation generating functional properties is a hallmark of living systems. Likewise, conductive protein assemblies switch between assembly states to regulate electrical responses through energy-consuming supramolecular processes. Supramolecular materials show promise in electronics as recoverable substrates in the future, exploiting multivalent non-covalent interactions. We sought to mimic dynamic supramolecular assemblies by modulating ATP concentrations in a controlled oscillatory manner using potato apyrase (PA) enzyme. PA is a phosphoesterase enzyme that hydrolyses ATP into AMP and inorganic phosphate (P_i_), thereby disrupting the ATP-templated assemblies.^44,45^ We hypothesized that PA would break down the nanowires, leading to decreased electrical response. Refuelling the system with additional ATP should reassemble the long nanowires with long-range charge transfer.

To begin with, we studied the morphology modulation of **Pt-1** and **Pt-1**.ATP using PA on a HOPG surface. AFM studies on these systems displayed the crosslinked **Pt-1**.ATP nanowires are broken down into small fragments and spherical structures, which may be due to the residual phosphate and monophosphate units (Figure 5a). Moreover, the PL intensity of the **Pt-1**.ATP assembly was reduced ∼14-fold to a value equivalent to that of **Pt-1** alone (Figure S36). Further, conductance measurements confirmed the breakdown of the supramolecular assembly of **Pt-1**.ATP. After PA treatment, the current density of the assembly showed a drastic reduction over two orders of magnitude aligning closely with the conductance level of the **Pt-1** complex alone (Figure 5b). This suggests that the phosphoesterase enzyme turns off the electrical transport across the nanowires by disrupting the efficient charge-conducting Pt···Pt interactions.

**Figure 5.**
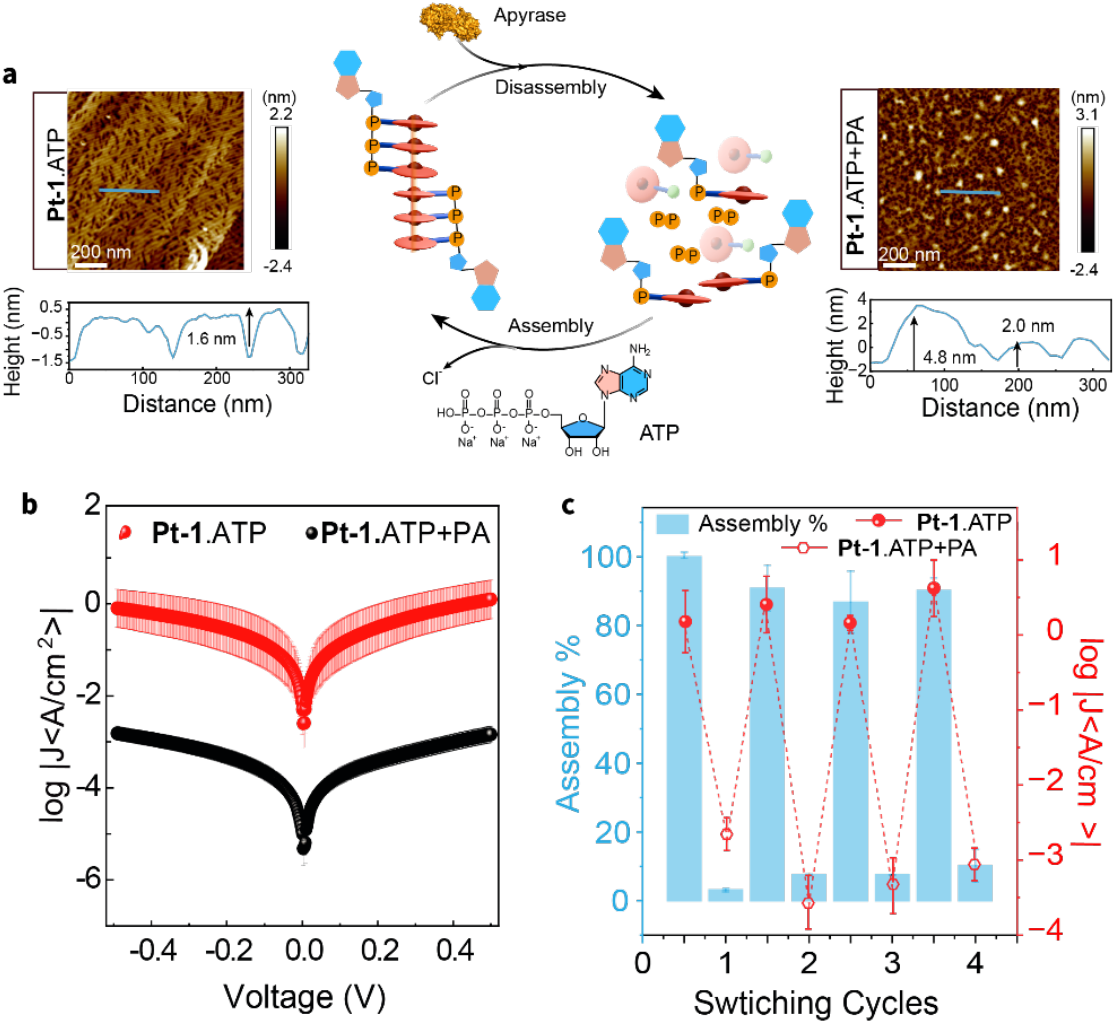
Enzymatic recycling of Pt-1.ATP supramolecular assemblies by potato apyrase. **a**, Schematic illustration of the reversible supramolecular assembly process. The **Pt-1** complex undergoes self-assembly into nanostructures upon binding with ATP, forming **Pt-1**.ATP assemblies. Upon enzymatic treatment with potato apyrase, ATP is hydrolyzed into AMP and inorganic phosphate, leading to disassembly of the supramolecular structure. Atomic force microscopy (AFM) image of **Pt-1**.ATP assembly showing extended nanowire-like structures formed upon ATP binding (left side). AFM image after treatment with potato apyrase, showing the breakdown of these assemblies and loss of ordered nanostructures (right side). **b**, Plot of log|J| versus voltage (V) for **Pt-1**.ATP (red) and after addition of potato apyrase (black), showing a significant decrease in current density upon enzymatic breakdown, consistent with the loss of conductive supramolecular architecture. **c**, Reversible switching behavior between high and low current density states (log|J| at –0.5 V) over multiple cycles, demonstrating dynamic control of the supramolecular assembly and disassembly through ATP binding and enzymatic hydrolysis.

Given the critical role of the supramolecular network in ensuring adaptability and recyclability, we sought to demonstrate multiple cycles of conductivity within our system to highlight its dynamic and reusable nature. Initially, **Pt-1**·ATP was spin-coated onto an HOPG substrate, and conductance measurements were performed under standard conditions. Subsequently, PA was introduced into the system by mixing it with ATP prior to spin-coating with **Pt-1**. This enzymatic intervention resulted in a significant drop in the current density of the assembly, bringing it to levels comparable to that of **Pt-1** alone. Replenishing ATP in the system remarkably restored the current density of the assembly to its initial level, highlighting the reversibility of its conductive behavior. To validate this reversible mechanism, we performed four consecutive cycles of assembly and disassembly, consistently observing similar results, as shown in Figure 5c and Figure S37-S38. This reproducibility underscores the robust and reversible nature of the **Pt-1**·ATP assembly’s conductance, which is intrinsically dependent on the presence of intact ATP within the supramolecular structure. The ability to repeatedly cycle between conductive and non-conductive states demonstrates the system’s dynamic adaptability. Such behaviour is promising for potential applications in energy storage, biosensing, or other ATP-responsive electronic devices.

## CONCLUSION

Drawing inspiration from biological systems and dissipative reaction networks, we have successfully developed a chemically driven transient assembly system. Herein, we focused on the use of triphosphate to facilitate the assembly of a benzimidazole-pyridine-based Pt(II) complex. The nucleophilic oxyanions in triphosphate effectively bind to the Pt(II) complex by substituting the chloro group, enhancing oligomerization through non-covalent interactions, particularly Pt···Pt and π-π interactions. This substitution results in the formation of longer and more stable supramolecular network that exhibits strong red luminescence due to metal-metal interactions. Notably, **Pt-1**.ATP showed more than a 4-fold increase in conductivity from ∼0.1 nS/cm to ∼1 µS/cm through electron transport to adjacent Pt–Pt centres.

A hallmark of living systems is their capability to dynamically switch between the active and inactive (dissipative) states. This switching is driven by the continuous presence of assembling components while executing essential biological functions. In this article, we showcase the oscillatory behavior of supramolecular oligomers through enzymatic ATP hydrolysis catalysed by potato apyrase. This enzymatic process ultimately leads to the depolymerization of the assembly. Our approach efficiently regulates both the Photoluminescence and conductivity. Overall, the study introduces a novel class of transient assemblies that draw inspiration from biological systems and possess advanced properties such as luminescence, conductivity, and temporal control. In future work, we will explore the potential applications of this system in fields such as biosensing, drug delivery, and intelligent chiral luminescent materials.

## Acknowledgements

S.R. acknowledges major financial support from Science and Engineering Research Board (CRG/2022/009021) and Department of Biotechnology (BT/PR49984/MED/32/911/2023). Seed grant by the IISc Digital Health Initiative is acknowledged for some consumables. V.K acknowledge the funding support from DST-INSPIRE (DST/INSPIRE/04/2018/002983), SERB-Core research grant (CRG/2020/002302). The support from Indian Institute of Science is gratefully acknowledged by SR and VK. The authors are thankful to the Department of Science and Technology (DST-FIST: SR/FST/PSII009/2010) for the instrumental facility at MRC. S.K. and R.K. acknowledge CSIR-UGC for the doctoral research fellowships. A.P. and P.S. are thankful to DST-Inspire and PMRF fellowship, respectively.

## Author contributions

S.K and R.K Contributed equally to this work.

